# The ascorbate-deficient guinea pig model of shigellosis allows the study of the entire *Shigella* life cycle

**DOI:** 10.1101/2020.08.28.270074

**Authors:** Antonin C André, Céline Mulet, Mark C Anderson, Louise Injarabian, Achim Buch, Verena Marina Prade, Axel Karl Walch, Jens Lykkesfeldt, Philippe Sansonetti, Benoit S Marteyn

## Abstract

*Shigella* spp. are the causative agents of bacillary dysentery or shigellosis, mainly in children living in developing countries. The study of *Shigella* entire life cycle *in vivo* and the evaluation of vaccine candidates’ protection efficacy have been hampered by the lack of a suitable animal model of infection (1). None of the ones evaluated so far (mouse, rabbit, guinea pig) allows to recapitulate shigellosis symptoms upon *Shigella* oral challenge. Historical reports suggest that dysentery and scurvy are both metabolic diseases associated with ascorbate-deficiency. Mammals which are susceptible to *Shigella* infection (humans, non-human primates and guinea pigs) are the lonely ones which are unable to synthesize ascorbate. We optimized a low-ascorbate diet to induce moderate ascorbate-deficiency but not scurvy in guinea pigs (Asc_plasma_ conc.=1.6 μM vs 36 μM with optimal ascorbate supply). We demonstrated that moderate ascorbate-deficiency increases shigellosis severity during extended period of time (up to 48h) with all strains tested (*Shigella flexneri* 5a and 2a, *Shigella sonnei*). At late time-points, a massive influx of neutrophils was observed both within the disrupted colonic mucosa and in the luminal compartment, although *Shigella* remains able to disseminate deep into the organ to reach the sub-mucosal layer and the bloodstream. This new model of shigellosis opens new doors for the study both of *Shigella* infection strategy and innate and adaptive immune responses to *Shigella* infection. It may be also of a great interest to study the virulence of other pathogen for which no suitable animal model of infection is available (*Vibrio cholerae, Yersinia pestis, Mycobacterium tuberculosis* or *Campylobacter jejuni*, among others).

**Significance:** The study of *Shigella* virulence cycle *in vivo* has been hampered by the lack of a suitable animal model, which would allow the colonic mucosa infection upon oral challenge. Based on historical reports and physiological aspects, it was suggested that ascorbate-deficiency may stand as a new dysentery risk-factor. To test this hypothesis, we set up a new ascorbate-deficient guinea pig model and demonstrated for the first time that the *Shigella* infectious process occurred for extended period of time (up to 48h) and demonstrated that shigellosis severity was higher in ascorbate-deficient animal. Ascorbate-deficient guinea pig model of infection may be used to assess the virulence of other pathogens for which no suitable animal model of infection is still lacking.

## Introduction

Bacillary dysentery or shigellosis remains nowadays a major burden disease especially in developing countries; annual shigellosis mortality was estimated in 2010 at 123,000 deaths worldwide among 88.4 million cases, mainly children under the age of five (2). In 2017, *Shigella* was included in the WHO list of the 12 antibiotic-resistant « priority pathogens » that pose the greatest threat to human health (3). Shigellosis is associated with fever, abdominal cramps and rectal inflammation. Dysenteric stools characteristically contain erythrocytes, polymorphonuclear neutrophils (PMNs) and mucus. Shigellosis is characterized by the specific invasion and destruction of the human colonic mucosa by *Shigella*, which is transmitted through the feco-oral route. No animal reservoir has been reported so far. Although the etiologic agents, *Shigella* spp., have been identified during the last century ago, shigellosis represents a major threat to public global health since no licensed vaccine is available.

Shigellosis main risk factors are the poor sanitation and the limited access to clean water (4). Malnutrition has been also shown to correlate with increased risk of shigellosis (5) (6). If no direct correlation between specific nutrient deficiency and shigellosis susceptibility has been reported, it has been shown that Vitamin A (7) or zinc supplementation (8) reduces the severity of acute shigellosis in malnourished children in Bangladesh and is recommended by the WHO as a supportive care in association with antimicrobial therapy (9).

In addition to these prophylactic treatments, the development of a *pan-Shigella* vaccine remains urgently needed. The lack of a suitable animal model of shigellosis - from an oral challenge to the specific infection and destruction to the colonic mucosa - represents a major drawback to evaluate vaccine candidates’ safety and efficacy (reviewed in (1)), but also to fully understand *Shigella* virulence cycle. Over the last decades several animal models of shigellosis have been used: monkeys, rabbits, mice, guinea pigs. All of them have their own limitation, none of them recapitulates all shigellosis symptoms (1). The infection of the colonic mucosa by *Shigella* following an oral challenge was only reported in rhesus macaques (*Macaca mulatta)* (10). A transient colonic infection has been reported in young guinea pigs (Hartley strain) upon intrarectal inoculation (11). In other proposed models, including the sereny test (guinea pig), the ileal loop model (rabbit) or the intravenous or intranasal infections (mouse), the targeted organs are different, making difficult the interpretation of the results. It is important to highlight here that upon oral challenge; mice are not susceptible to *Shigella* infection for reasons which remain elusive.

An efficient immunological response to *Shigella* infection (including IgA and IgG secretion) has been observed in human and in several animal models (monkeys, mice, rabbit, guinea pigs). It has been reported that this response is serotype-specific and mainly directed toward *Shigella* lipopolysaccharide (LPS-associated O-antigen) (12). However, no direct correlation between the induced immune response and a potential protection against has been confirmed. Currently, vaccine candidates are evaluated in clinical trials (phases I-III) only based on their immunogenicity assessed in preclinical models (mainly mice), with no proof of efficacy against *Shigella* infection. This strategy is time-consuming, risky and expensive and should be optimized to improve chances of success (13); this includes the validation of a suitable animal model of shigellosis.

In this study, we aimed at identifying novel shigellosis susceptibility factors to develop and validate an animal model of *Shigella* which would faithfully reproduce all shigellosis physiopathology and clinical symptoms. We hypothesized that an ascorbate-deficiency may be an unrevealed shigellosis susceptibility risk factor, based on physiological bases and historical reports.

First, acute shigellosis has been reported so far only in humans, non-human primates and guinea pigs, which are the lonely mammal species unable to synthesize ascorbate, due to the deficiency in expressing a functional L-gulonolactone oxidase enzyme (GULO), converting L-gulono-1,4-lactone in L-ascorbic acid (ascorbate or Vitamin C). Conversely, mice express a functional GULO synthesize ascorbate and do not develop diarrhea upon *Shigella* infection (14), suggesting that ascorbate optimal supply may limit shigellosis severity. Conversely, Honjo and colleagues have shown that shigellosis symptoms were more severe in ascorbate-deficient monkeys (cynomolgus monkeys) upon oral challenge (15).

Second, the association of scurvy - caused by severe ascorbate deficiency - and dysentery has been first reported by Lind in 1753 (16). Since then, it was hypothesized that both scurvy and chronic diarrhea may be nutritional deficiency syndromes, affecting populations suffering from malnutrition (ie. soldiers, seamen) (17). Diarrhea was considered as a *“symptom”* or *“part”* of scurvy and characterized as *“scorbutic fluxes”* or *“scorbuticdiarrhea”.* Importantly, ascorbate supplementation allowed the treatment of both scurvy and diarrhea symptoms (16). Since these preliminary observations, if severe ascorbate-deficiency has been confirmed as the main cause of scurvy, no clinical symptom or diseases have been associated with moderate ascorbate-deficiency.

In this report, we developed and validated an ascorbate-deficient guinea pig model of shigellosis, which recapitulates the shigellosis symptoms upon oral challenge with *Shigella flexneri* (*S. flexneri*) 2a and 5a, and *Shigella sonnei* (*S. sonnei*).

## Results

### Optimization of the guinea pig diet to induce a moderate ascorbate-deficiency without scurvy symptoms

The guinea pig model of shigellosis reporting the colonization and destruction of the colonic mucosa by *Shigella* has been described by Shim and colleagues (11) and consists in an intrarectal challenge of young guinea pigs (2-week) with *S. flexneri* 2a and *S. flexneri* 5a. It was reported that a severe and acute colitis is induced 8h post-infection (p.i.) and that symptoms disappear by 24h-48h p.i., thus representing an important limitation of the model. In addition, it was shown that older animals (5-week and more) are no longer susceptible to *Shigella* infection (11). We confirmed these results in our laboratory and used this model in routine in various studies to characterize *Shigella* virulence mechanisms *in vivo* (18–20).

In this study, we aimed at evaluating the impact on shigellosis severity of a moderate ascorbate-deficiency in guinea pigs. In order to proceed, we fed animal for 2 weeks with specific diets containing sub-optimal ascorbate amounts (Fig. 1A). The amount of ascorbate in diet allowing an optimal supply to guinea pigs is 400 mg ascorbate/kg (Safediet, see Methods). We evaluated the impact of diets containing 50, <50 and 0 mg ascorbate/kg (Fig. 1A) on guinea pig plasma ascorbate concentration (Fig. 1B), animal weight (Fig. 1C and S1) and reported the absence of scurvy symptoms (Fig. 1D).

**Figure 1.**
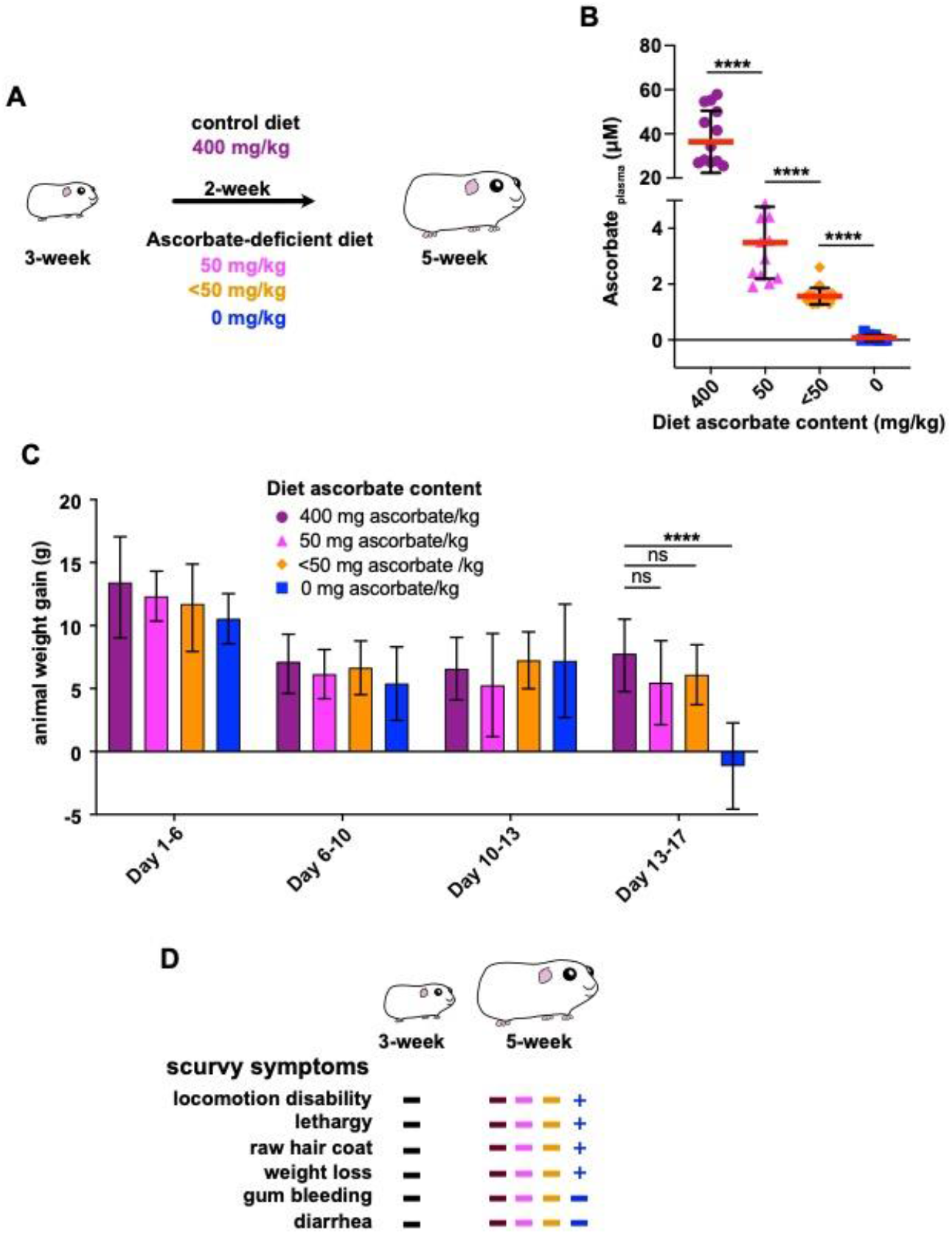
Optimization of diet ascorbate content to induce guinea pig moderate ascorbate-deficiency. (**A**) Young guinea pigs (3-week) were fed for two weeks with diets containing various ascorbate concentration to induce moderate ascorbate-deficiency without inducing scurvy, caused by severe ascorbate deficiency. Commercial diet contains 400 mg ascorbate/kg (Safediet); suboptimal ascorbate supply was obtained with diets containing 50 mg, <50 mg and 0 mg ascorbate/kg. (**B**) Guinea pig ascorbate deficiency was confirmed by dosing plasma ascorbate concentration. Results are expressed as Mean ± S.D., **** indicates T-test *p*<0.0001 (n>10 animals per group). (**C-D**) Moderate ascorbate deficiency induction do not lead to significant reduction of weight gain (see also Fig. S1), animals were weighted regularly (Day 6, 10, 10, 13 and 17) during this period of time. Results are expressed as Mean ± S.D., ‘ns’ indicates T-test *p*>0.05, **** indicates *p*<0.0001 (n>10 animals per group). (**D**) The absence of scurvy symptoms (as indicated) was confirmed in young animals (3-week) and after 2-week feeding with diets containing 400, 50, <50 and 0 mg ascorbate/kg. ‘+’ indicates the corresponding scurvy symptom has been observed in at least one animal of the group.

The guinea pig plasma ascorbate concentration upon optimal ascorbate supply is 36.4 ± 14.1 μM, confirming results from other studies (21), which is in the same range although lower compared to human plasma ascorbate concentration (49.5 ± 14.2 μM (22)). When guinea pig diet is devoid of ascorbate (0 mg/kg), the plasma ascorbate concentration is 0.1 ± 0.1 μM, as anticipated, confirming that a 2-week diet is sufficient to induce ascorbate-deficiency. A moderate ascorbate-deficiency is induced when guinea pigs are fed with a diet containing 50 mg ascorbate/kg (3.5 ± 1.3 μM) or <50 mg/kg (1.6 ± 0.3 μM) (Fig. 1B). We further aimed at defining the most suitable low-ascorbate diet, which ensure a significant reduction of guinea pig plasma ascorbate concentration while preventing scurvy.

Animal weight of all groups increased for 13 days and were not significantly different at each timepoint (Fig. S1, *p*>0.05). However, between Day 13 and Day 17, a significant weight loss has been observed in the absence of ascorbate supply (Fig. 1C, 0 mg ascorbate/kg, *p*<0.001), not with diets containing sub-optimal ascorbate amounts (Fig. 1C, *p*>0.05). In addition, no scurvy symptom (locomotion disability, lethargy, raw hair coat or weight loss) was observed in animal fed with this diet, as opposed to the one devoid of ascorbate (Fig. 1D).

In conclusion, a diet containing <50 mg ascorbate/kg induces a significant plasma ascorbate concentration decrease (Fig. 1B), without affecting animal growth (Fig. 1C and S1) or causing scurvy (Fig. 1D) and will further be used in this study.

### Moderate ascorbate-deficiency increases shigellosis severity upon *Shigella* intrarectal challenge

In human, shigellosis symptoms appear 24h to 48h p.i.. As mentioned above, guinea pigs fed with an optimal diet do no develop such long-term infectious processes (*Shigella* being cleared by the immune system 24h p.i.). In addition, only young guinea pigs (3-week, just after weaning) are susceptible to *Shigella* infection inoculated at high-dose intrarectally (11): older animals (5-week or over) are not. An oral *Shigella* challenge do not lead to the infection and destruction of the colonic mucosa, as occurring in humans (fecal-oral route).

Using *S. flexneri* 5a as a model, we first confirmed that the colonic mucosa is disrupted 8h p.i. by *Shigella* upon intrarectal challenge in young animals (3-week) fed with a high-ascorbate diet (400 mg ascorbate/kg) (Fig. 2). We confirmed that older animals (5-week) fed with a similar diet were not susceptible to *Shigella* infection (Fig. 2). Conversely, we observed that ascorbate-deficient guinea pigs (5-week) remained susceptible to *S. flexneri* 5a infection (colonic mucosa destruction, leucocyte infiltration, oedema); this phenotype was observed in animals fed with both low-ascorbate diets (50 and <5à mg ascorbate/kg) (Fig. 2). However not significant weight loss was observed 8h p.i. in 5-week guinea pigs (due to high variability of the results), as observed in young animals (Fig. S2, *p*<0.05) and previously reported (11).

**Figure 2.**
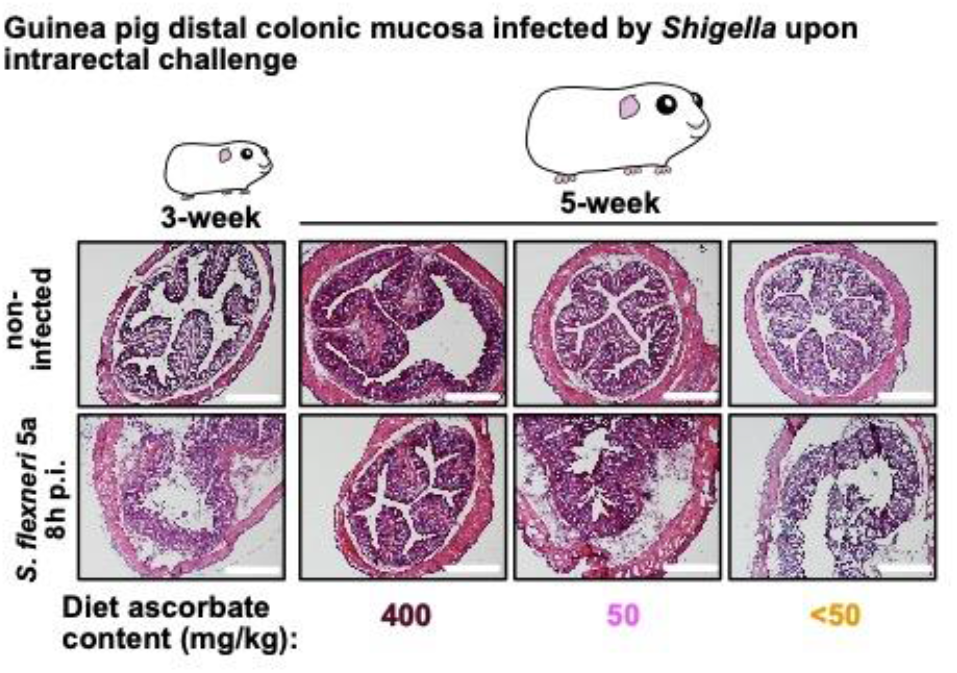
Moderate ascorbate-deficiency increases the susceptibility of older guinea pigs to *Shigella* infection upon intrarectal challenge. Young (3-week) and older guinea pigs fed with 400, 50 and <50 mg ascorbate/kg diets (5-week) were challenged intrarectally with 10_10_ c.f.u. *S. flexneri* 5a (M90T). 8h p.i. animals were sacrificed, and distal colonic samples were collected, stained with haematoxylin eosin and compared to non-infected tissues. Scale bars are 30 mm. Animal weight was measured before and after infection; these results are shown in Fig. S2.

We hypothesized that weight-loss may be observed in 5-week guinea pigs during long-term infections. To address this question, the Shigella infection course was studied in ascorbate-deficient guinea pigs for longer infection period of time using various *Shigella* species.

### Prolonged infection processes are induced by *S. flexneri* 5a, *S. flexneri* 2a and *S. sonnei* in ascorbate-deficient guinea pigs

We evaluated the severity of *Shigella* infection 30h p.i. upon intrrectal infection in 5-week ascorbate-deficient guinea pigs, as compared to control (Fig. S3A). We first confirmed that no difference in animal weight was observed during the period of ascorbate deficiency induction (Fig. S3A, *p*>0.05).

As previously reported (11), we did not observe severe symptoms of shigellosis 30h post-infection in guinea pigs fed with a 400 mg ascorbate/kg diet (Fig. S3B) Infected animals kept gaining after during infection period (30h, p>0.05), also in a lesser extent compared to non-infected animals (p<0.01) (Fig. S3B), confirming that a transitory infection occurred at early time-point (8h p.i., Fig. 2).

Conversely, severe *Shigella* infections were observed 30h p.i. (intrarectal challenge) in 5-week ascorbate-deficient guinea pigs (colonic mucosa destruction, leucocyte infiltration, oedema), using *S. flexneri* 5a, *S. flexneri* 2a or *S. sonnei*, not with an avirulent plasmid-cured *S. flexneri* 5a *ΔmxiD* mutant (Fig. 3A). If non-infected guinea pigs kept gaining weight during this period of time, a significant weight loss is induced by *Shigella* infection in all groups (Fig. 3B, *p*<0.01 or *p*<0.05), not with *S. flexneri* 5a *ΔmxiD* (Fig. 3B, *p*>0.05). These results demonstrate that ascorbate-deficiency increases the severity of *Shigella* spp. infection in guinea pigs. We previously reported that Shigella infection induces a significant decrease of plasma ascorbate concentration (22), through a mechanism which remains to be defined. Here, we further investigated in a complementary approach the abundance of ascorbate within the different colonic mucosa layers and its modulation during *Shigella* infection.

**Figure 3.**
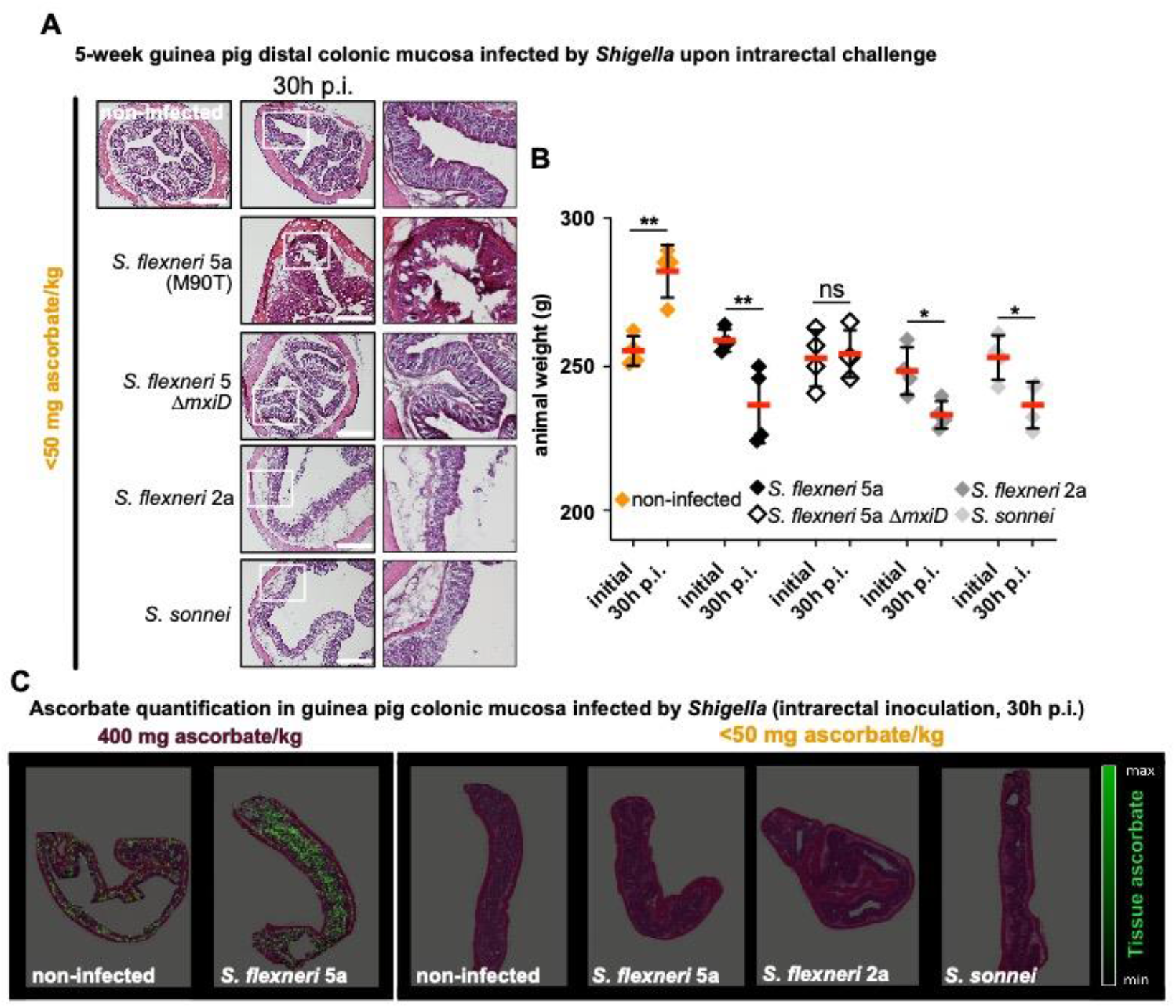
*Shigella* spp. colonize the colonic mucosa of ascorbate-deficient guinea pigs for extended period of time; ascorbate-deficiency is associated with a local reduction of ascorbate within colonic mucosa layers. (**A**) Ascorbate-deficient guinea pigs (<50 mg ascorbate/kg diet) were infected intrarectally with *S. flexneri* 5a (wt and *ΔmxiD* strains), *S. flexneri* 2a and *S. sonnei.* 30h p.i. animals were sacrificed, and distal colonic samples were collected, stained with haematoxylin eosin and compared to non-infected tissues. Scale bars are 30 mm. On the right panel, x3 magnification of white squares (left panel) are shown. (see also Fig. S3B for control animals fed with a high-ascorbate diet) (**B**) Animals were weighted before infection and 30h p.i. (see also Fig. S3C for control animals fed with a high-ascorbate diet). Results are expressed as Mean ± S.D., ‘ns’ indicates T-test *p*>0.05, * indicates *p*<0.05, ** indicates *p*<0.01 (4 animals per group). (**C**) Ascorbate abundance was quantified within the different layers of the colonic mucosa by imaging mass spectrometry in tissues from animals fed with 400 mg ascorbate/kg and <50 mg ascorbate/kg diets, infected or not with indicated strains for 30h (intrarectal challenge). Quantification in different layers are shown in Fig. S4B.

### *Shigella* infection induces a systemic ascorbate concentration decrease and occurs in low-ascorbate microenvironments

First, we confirmed in control animals (fed with high-ascorbate diet) the decrease of the plasma ascorbate concentration upon *S. flexneri* 5a infection (30h p.i., intrarectal challenge) (Fig. S4A, *p*<0.05). A significant decrease was observed in ascorbate-deficient guinea pigs upon *S. flexneri* 5a infection (30h p.i., intrarectal challenge) but not with *S. flexneri* 5a *ΔmxiD* mutant; despite the fact that the plasma ascorbate concentration was already low in basal conditions (Fig. S4A, *p*<0.01).

Then we quantified ascorbate abundance within the colonic mucosa layers by mass imaging spectrometry (see Methods) in the same models of *Shigella* infection (Fig. 3C and S4B). The determination of local ascorbate abundance remains difficult and only few quantitative data are available in colon (23).

We first observed that in basal conditions the ascorbate distribution is heterogeneous within the colon layers. In control animals (fed with high-ascorbate diet), higher amounts of ascorbate were detected within the submucosa and the muscularis mucosa, as compared to the mucosa or the epithelium (Fig. S4B, *p*<0.05 and *p*<0.01). In ascorbate-deficient guinea pigs, ascorbate was almost no detected in any layer under basal conditions (Fig. 3C (right panel) and S4B), showing for the first time that when ascorbate-deficiency is induced, not only the plasma ascorbate concentration but also its local distribution within the colon is strongly decreased. No impact of *Shigella* infection on ascorbate abundance within the colonic layers was observed in control or ascorbate-deficient animals (Fig. 3C and 4B, *p*>0.05). However, ascorbate is not detected within foci of infection both in control and ascorbate-deficient animals (Fig. S4B).

**Figure 4.**
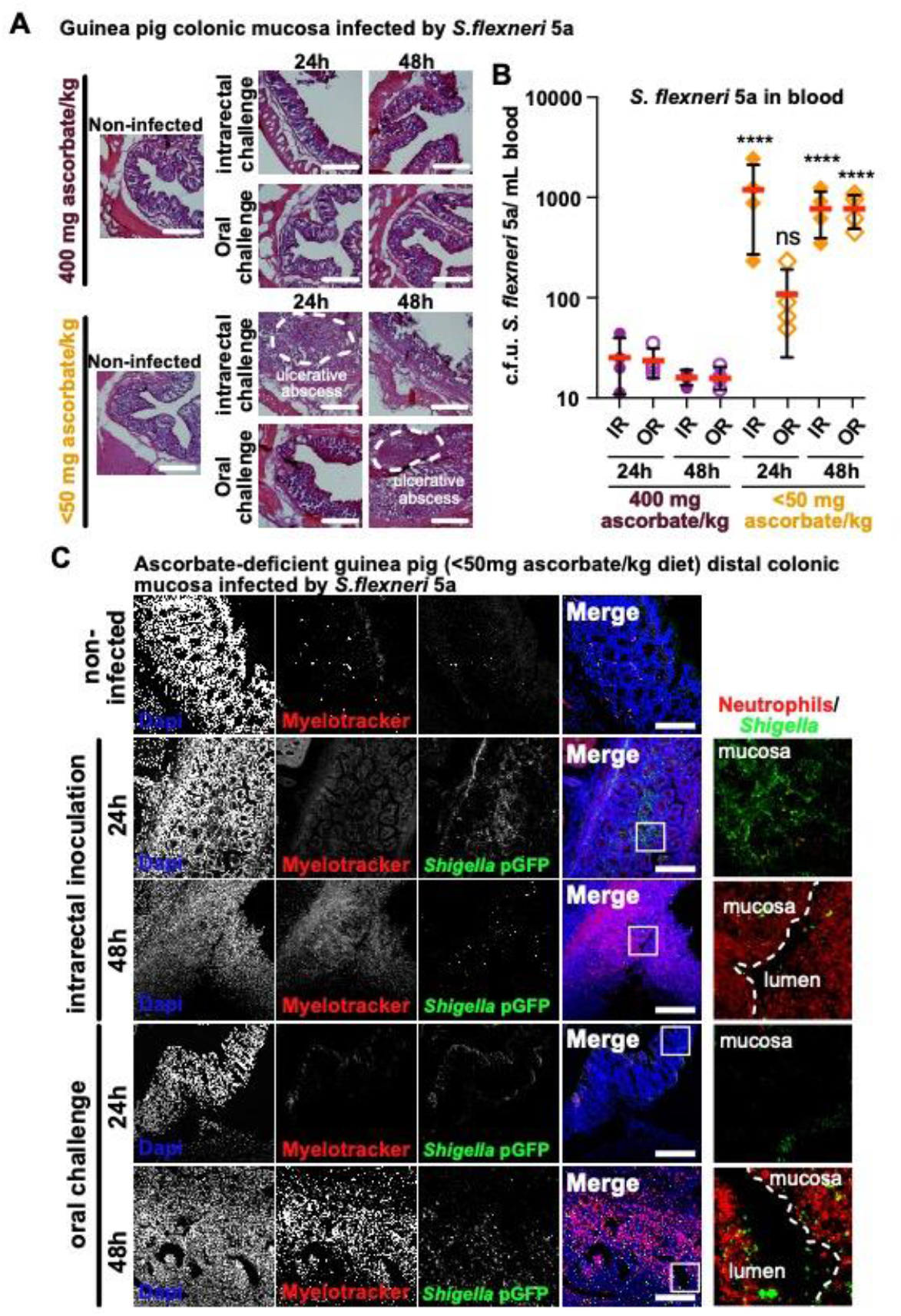
Ascorbate-deficient guinea pigs are susceptible to *Shigella* infection, upon intrarectal or oral challenge, over extended periods of time. (**A**) Ascorbate-deficient guinea pigs (<50 mg ascorbate/kg diet) and control animals (400 mg ascorbate/kg diet) were infected intrarectally or orally with 1010 c.f.u. *S. flexneri* 5a. 24h or 48h p.i. animals were sacrificed, and distal colonic samples were collected, stained with haematoxylin eosin and compared to non-infected tissues. Scale bars are 400 μm. (**B**) *S.flexneri* 5a potential translocation to the bloodstream was assessed by plating blood samples on TSA culture plates. Results are expressed as Mean ± S.D., ‘ns’ indicates T-test *p*>0.05, **** indicates *p*<0.0001 (4 animals per group). (**C**) *S. flexneri* 5a pGFP (green) and neutrophils were detected by immunofluorescence within the distal colonic mucosa of ascorbate-deficient guinea pigs (<50 mg ascorbate/kg diet) challenged orally or intrarectally for 24 and 48h with 1010 c.f.u.. DNA was stained with Dapi (blue), neutrophils were stained with Myelotracker-Cy3 (18) (red). Scale bars are 150 μm. On the right panel, x5 magnification of white squares (left panel) are shown. Detection of *S. flexneri* pGFP and neutrophils in tissues from control animals (400 mg ascorbate/kg diet) is shown in Fig. S6.

Since we confirmed that ascorbate-deficiency increase shigellosis symptom severity, we further aimed at evaluating whether the overall life cycle of *Shigella* (from an oral challenge to the colon colonization) could be recapitulated in this new shigellosis animal model during extended periods of infection.

### Efficient colonization of the colonic mucosa by *Shigella* is observed in ascorbate-deficient guinea pigs

To recapitulate *Shigella* life cycle, control and ascorbate-deficient guinea pigs were challenged orally with *S. flexneri* 5a pGFP and long-term infections (24h and 48h) were performed (Fig. S5A): animals were weighted and the colonic mucosa colonization and destruction by *Shigella* was then evaluated. As a major sign of shigellosis severity, we demonstrated that ascorbate-deficient animal weight continuously decreased during the time of infection, reaching 20% weight-loss 48h p.i. upon oral and intrarectal challenge (Fig. S5B, *p*<0.05 and *p*<0.01 respectively); this phenotype was not observed in control animals (Fig. S5B). We observed comparable inflammation and infection symptoms within the colonic mucosa at 24h and 48h p.i. upon intrarectal and oral challenge major : oedema were observed until late timepoints (48h p.i.) and the formation of large and purulent ulcerative abscesses were formed with the colonic submucosa, in association with the destruction of the tissue (Fig. 4A). No sign of tissue destruction or inflammation was observed in control animals (fed with high-ascorbate diet) using similar experimental procedures (Fig. 4A). In addition, we have shown that *Shigella* could reach the blood circulation in ascorbate-deficient animals (oral and intrarectal challenge), not in control animals (Fig. 4B, *p*<0.0001). To investigate in more details the *Shigella* infectious process we performed an immunolabeling of *Shigella* and neutrophils and observed that *Shigella* colonized deep colonic layers such as the submuscularis mucosa and the mucosa of ascorbate-deficient animals. A massive recruitment of neutrophils was additionally observed within the infected tissues 24h p.i., without impairing *Shigella* dissemination 48h p.i. (Fig. 4C). In addition, large populations of both *Shigella* and neutrophils were detected within the luminal compartment, as reported in patients suffering of shigellosis (Fig. 5C), confirming the efficient dissemination of the bacteria during long-term infection. Conversely, in control animals, no destruction of the tissue was observed 24h or 48h p.i., with limited colonization of the mucosa by *Shigella* and only few neutrophils were recruited (Fig. S6).

These results confirmed that ascorbate-deficiency increases shigellosis symptom severity during extended periods of infection; this animal model of shigellosis allows for the first time to follow *Shigella* life cycle from oral challenge to the colonization and destruction of the colonic mucosa, despite of a the innate immune response induction, as reported in humans.

## Discussion

While severe ascorbate-deficiency is causing scurvy, we demonstrate here that moderate ascorbate-deficiency is new shigellosis risk-factor, which has been so far largely underestimated. This new insight raises the question of the definition of a moderate ascorbate deficiency, which is currently not well characterized in humans, probably due to the difficulty to dose plasma ascorbate in routine. In guinea pigs, our results show that plasma ascorbate concentrations from 1.6 ± 0.3 μM to 3.5 ± 1.3 μM (vs 36.4 μM in animals fed with a high-ascorbate diet) are associated with increased shigellosis severity (Fig. 1A and Fig. 2). Further studies are required to define upper values of moderate ascorbate-deficiency defined in this study. In humans, optimal ascorbate plasma concentration is estimated around 50 μM (confirmed in our previous work (49.5 ± 14.2 μM) (22)), but no assessment plasma ascorbate concentrations associated with moderate deficiency has been achieved so far and requires further clinical studies, especially in population suffering of shigellosis (mainly children under the age of 5, in developing countries). In a previous study it was reported that 20% of children in Puerto Rico were moderately deficient (<30 μM), with no correlation with diarrhea episode (24). These clinical data will be of a great interest to confirm the relevance and potential optimization of the animal model of shigellosis characterized in this study. Until now, epidemiological data are lacking in resource-limited countries (RLC), to correlate ascorbate-deficiency and diarrhea episodes determine whether ascorbate deficiency is especially in children suffering from diarrhea.

We have validated here a new animal model of shigellosis, the ascorbate-deficient guinea pig, which allows to follow for the first time the overall *Shigella* life cycle from an oral challenge to the colonization and destruction of the colonic mucosa (Fig. 4A-C). Until now, only transient infections were observed, associated with foci of infection formation within the upper part of the colonic submucosa (20, 25), *Shigella* being further cleared during the immune response in this model. The ascorbate-deficient guinea pig model of shigellosis will be of a great help to investigate the role of the different immunoglubulins and immune cell populations to limit the dissemination of *Shigella* within the colonic mucosa, but also, conversely, to better understand the molecular bases of the subversion and their antimicrobial activity by *Shigella* to mediates its efficient dissemination within the infected tissues and to the bloodstream as reported here (Fig. 4B). More specifically, this new shigellosis model will allow to take a fresh look at the role of mucosal IgA in preventing Shigella dissemination(26, 27) and at *Shigella* capacity to subvert immune-cell function and anti-microbial activity, which has been described previously *in vitro* or *in vivo* in other models (macrophage and B-lymphocyte apoptosis induction (28, 29) or T-cell migration (30, 31)). Most of these subversion mechanisms are dependent on *Shigella* Type Three Secretion System (T33S). Since we have shown in a previous study that this secretion apparatus is mainly inactive within hypoxic foci of infection (20), alternative virulence mechanisms are expected to be involved and may be identified in this new model of shigellosis.

Last, the ascorbate-deficient guinea pig model of shigellosis described in this study will allow to investigate the composition, structural organization and role of Isolated Lymphoïd Follicles (belonging to the Gut-associated Lymphoid Tissue (GALT), specifically disseminated within the colonic mucosa) during *Shigella* infection. which have not yet been characterized.

## Materials and Methods

### Bacteria

Unless otherwise noted, *S. flexneri* 5a (M90T, wt and *ΔmxiD* mutant) and 2a strains were grown at 37°C in Tryptic Soy Broth (TSB) with shaking or TSB agar plates supplemented with 0.01% Congo Red (Sigma-Aldrich) and Ampicillin (100 μg/ml) when bacteria were transformed with the pGFP plasmid. *S. sonnei* was acquired from the Institut Pasteur strain collection (CIP 106347) and is a clinical isolate from a 1999 Paris infection (32). The strain was grown in TSB supplemented with Ampicillin (100 μg/ml) to maintain the pMW211 plasmid.

### Ascorbate-deficient guinea pig model of shigellosis

Young conventional guinea pigs (3 weeks old; female; Dunkin–Hartley; <150 g) were used to study *Shigella* infection severity. Guinea pigs were fed for fifteen days with a standard diet allowing an optimal ascorbate supply (400 mg ascorbate/kg, Safediet ref. 106) as previously described (33) or an ascorbate-deficient diets specifically designed and produced by Safediet (0, 10 or 50 mg ascorbate/kg). Ascorbate-deficiency was assessed by dosage of its plasma concentration (see below). Guinea pig infection with *Shigella* infection was achieved with *Shigella flexneri* 5a wild-type and *ΔmxiD* strains, *Shigella flexneri* 2a and *S. sonnei* wild-type strains was performed by intrarectal challenge (as described previously (20)) or oral challenge of animals with 10_10_ c.f.u. exponentially grown, as indicated. Infections proceeded for 8 h, 24h, 30h or 48h, as indicated, before animals were sacrificed. Experimental procedures were approved by the Institut Pasteur ethics committee (auth. n°190127).

The distal colon was sectioned (7 cm) and either flash-frozen in liquid nitrogen for tissue ascorbate abundance quantification (imaging mass spectrometry, see below) or fixed in 4% PFA in 1× PBS for 1–2 h and then incubated in 1× PBS/glycine (100 mM) for 30 min to quench the PFA (for tissue labeling and imaging). PFA-fixed tissues were immersed successively in 15% and 30% sucrose at 4 °C overnight. Tissue transversal sections (1 cm) were embedded in Tissue-Tek O.C.T. compound (Sakura) using a flash-freeze protocol and frozen at −80 °C. For histological and standard immunofluorescence staining, 10-μm sections were obtained using a CM-3050 cryostat (Leica Biosystems).

To quantify the abundance of bacteria in the bloodstream, 100 μL blood samples were collected in the presence of EDTA and plates on GTCS + Ampicillin agar plates. Bacteria were counted after overnight culture at 37°C.

### Plasma ascorbate dosage

Guinea pig blood samples were collected by intracardiac puncture in the presence of EDTA. Following centrifugation for 5 min at 2,000 *x g*, the plasma was acidified with an equal volume of 10% (w/v) metaphosphoric acid (MPA) containing 2 mmol/L of disodium-EDTA. Ascorbate concentration was quantified by high-performance liquid chromatography with coulometric detection, as described previously (22, 34). Results were averaged from dosage performed on at least 4 animals per conditions.

### Tissue ascorbate quantification

Imaging mass spectrometry was performed in collaboration with the team of Axel Walch (Helmholtz Zentrum München) on flash-frozen colonic samples.

Metabolite mapping was done using frozen (12 μm, Leica Microsystems, CM1950, Germany) guinea pig colonic tissue samples, mounted onto indium-tinoxide (ITO)-coated glass slides (Bruker Daltonik, Bremen, Germany) pretreated with 1:1 poly-Llysine (Sigma Aldrich, Munich, Germany) and 0.1% Nonidet P-40 (Sigma). The air-dried tissue sections were spray-coated with 10 mg/ml 9-aminoacridine hydrochloride monohydrate matrix (Sigma-Aldrich, Munich, Germany) in 70% methanol using the SunCollect™ sprayer (Sunchrom, Friedrichsdorf, Germany). Spray-coating of the matrix was conducted in eight passes (ascending flow rates 10 μl/min, 20 μl/min, 30 μl/min for layers 1–3, and layers 4–8 with 40 μl/min), utilizing 2 mm line distance, and a spray velocity of 900 mm/min. Metabolites were detected in negative-ion mode on a 7T Solarix XR Fourier-transform ion cyclotron resonance (FTICR) mass spectrometer (Bruker Daltonik) equipped with a dual ESI-MALDI source and a smartbeam-II Nd:YAG (355 nm) laser. Data acquisition parameters were specified in ftmsControl software 2.2 and flexImaging (v. 5.0) (Bruker Daltonik). Mass spectra were acquired in negative-ion mode covering /75–1100, with a 1M transient (0.367 sec duration), and an estimated resolving power of 49,000 at m/z 200,000. The laser operated at a frequency of 1,000 Hz utilizing 200 laser shots per pixel with a pixel resolution of 15 μm. L-Arginine was used for external calibration in the ESI mode. On-tissue MS/MS was conducted on guinea pig colonic tissue section using continuous accumulation of selected ions’ mode and collision-induced dissociation (CID) in the collision cell. MS/MS spectra were analyzed by Bruker Compass DataAnalysis 5.0 (Build 203.2.3586).

### Tissue labeling and imaging

For histological studies, sectioned colonic samples were stained with hematoxylin-eosin following standard procedures (described in (35) Labeled tissues were imaged with a transmitted light microscope (Axioskop 2, Zeiss) using x10 or x20 objectives.

For immunofluorescence imaging, PFA-fixed sectioned tissues were washed three times in PBS + 0.1% saponin prior immunolabeling in the same buffer for 1 hour. Nuclei were stained with Dapi (1 mg/mL, Life Technologies) (1:1000) and Myelotracker-Cy3 (1 mg/mL, 1:1000), a specific neutrophil marker designed and validated in our laboratory (18). After three additional washes in PBS + 0.1% saponin, three times in PBS and three times in deionized H20 and mounted with Prolong gold mounting media (Thermofisher Scientific). Immunolabeled guinea pig colons infected with *Shigella* spp. pGFP strains were imaged on a laser-scanning TCS SP5 confocal microscope (Leica). Z-stack images were taken with 1 μM step-size increments. Obtained Z-stack images were processed with Fiji software

### Statistics

Data were analyzed with the Prism 8 software (GraphPad). ANOVA or Student T-test were performed to analyze the different datasets.

## Supporting information

Supplementary informations

## Acknowledgments

We thank Patricia Flamant from the Institut Pasteur Histology Platform for processing guinea pig colonic tissues. We are very grateful to team of the Institut Pasteur Animalerie Centrale for their constant support to this project (particularly Serge Hedan, Marion Berard and Myriam Mattei). This work was supported by the ANR JCJC grant (ANR-17-CE15-0012) (B.S.M.).

## References

1. M. BS, Shigella Vaccine Development: The Model Matters. JSM Trop Med Res 2, 1011–1014 (2016).

2. R. Lozano, et al., Global and regional mortality from 235 causes of death for 20 age groups in 1990 and 2010: a systematic analysis for the Global Burden of Disease Study 2010. Lancet 380, 2095–2128 (2012).

3. E. Tacconelli, et al., Discovery, research, and development of new antibiotics: the WHO priority list of antibiotic-resistant bacteria and tuberculosis. Lancet Infect Dis 18, 318–327 (2017).

4. Water and Sanitation-Related Diseases and the Environment: Challenges, Interventions, and Preventive Measures. Edited by Janine M. H. Selendy. Hoboken (New Jersey): Wiley-Blackwell. $139.95 (paper). xvii + 497 p. + 16 pl.; ill.; index. ISBN: 978-0-470-52785-6. [Two DVDs are included.] 2011. Q Rev Biology 88, 238–238 (2013).

5. M. M. Mahbub, C. R. Ahsan, M. Yasmin, J. Nessa, Analysis of Different Prognostic Indicators for Malnutrition and Shigella flexneri Infection Among the Children in Bangladesh. Indian J Microbiol 52, 400–5 (2012).

6. F. Ferdous, et al., Severity of diarrhea and malnutrition among under five-year-old children in rural Bangladesh. Am J Tropical Medicine Hyg 89, 223–8 (2013).

7. S. Hossain, et al., Single dose vitamin A treatment in acute shigellosis in Bangladeshi children: randomised double blind controlled trial. Bmj 316, 422–426 (1998).

8. S. K. Roy, et al., Zinc supplementation in the management of shigellosis in malnourished children in Bangladesh. Eur J Clin Nutr 62, 849–855 (2008).

9. WHO, “Guidelines for the control of shigellosis, includingepidemics due to Shigella dysenteriae type 1” (2005).

10. A. Fontaine, J. Arondel, P. J. Sansonetti, Role of Shiga toxin in the pathogenesis of bacillary dysentery, studied by using a Tox-mutant of Shigella dysenteriae 1. Infect Immun 56, 3099–109 (1988).

11. D.-H. Shim, et al., New Animal Model of Shigellosis in the Guinea Pig: Its Usefulness for Protective Efficacy Studies. J Immunol 178, 2476–2482 (2007).

12. M. M. Levine, K. L. Kotloff, E. M. Barry, M. F. Pasetti, M. B. Sztein, Clinical trials of Shigella vaccines: two steps forward and one step back on a long, hard road. Nat Rev Microbiol 5, 540–553 (2007).

13. H. Wenzel, et al., Improving chances for successful clinical outcomes with better preclinical models. Vaccine 35, 6798–6802 (2017).

14. M. A. McArthur, M. Maciel, M. F. Pasetti, Human immune responses against Shigella and enterotoxigenic E. coli: Current advances and the path forward. Vaccine 35, 6803–6806 (2017).

15. S. Honjo, M. Takasaka, T. Fujiwara, K. Imaizumi, H. Ogawa, SHIGELLOSIS IN CYNOMOLGUS MONKEYS (MACACA IRUS). Jpn J Medical Sci Biology 22, 149–162 (1969).

16. J. Lind, A Treatise of the Scurvy in Three Parts (1753).

17. A. J. Bollet, Scurvy and chronic diarrhea in Civil War troops: were they both nutritional deficiency syndromes? J Hist Med All Sci 47, 49–67 (1992).

18. M. C. Anderson, et al., MUB40 Binds to Lactoferrin and Stands as a Specific Neutrophil Marker. Cell Chem Biol 25, 483–493.e9 (2018).

19. B. S. Marteyn, et al., ZapE Is a Novel Cell Division Protein Interacting with FtsZ and Modulating the Z-Ring Dynamics. Mbio 5, e00022–14 (2014).

20. J.-Y. Tinevez, et al., Shigella-mediated oxygen depletion is essential for intestinal mucosa colonization. Nat Microbiol 4, 2001–2009 (2019).

21. A. Mortensen, S. Hasselholt, P. Tveden-Nyborg, J. Lykkesfeldt, Guinea pig ascorbate status predicts tetrahydrobiopterin plasma concentration and oxidation ratio in vivo. Nutrition Res New York N Y 33, 859–67 (2013).

22. L. Injarabian, et al., Ascorbate maintains a low plasma oxygen level. Sci Rep-uk 10, 10659 (2020).

23. S. J. Padayatty, M. Levine, Vitamin C: the known and the unknown and Goldilocks. Oral Dis 22, 463–93 (2016).

24. A. M. Preston, C. Rodríguez, C. E. Rivera, Plasma ascorbate in a population of children: influence of age, gender, vitamin C intake, BMI and smoke exposure. P R Health Sci J 25, 137–42 (2006).

25. E. T. Arena, et al., Bioimage analysis of Shigella infection reveals targeting of colonic crypts. Proc National Acad Sci 112, E3282–E3290 (2015).

26. A. Phalipon, et al., Monoclonal immunoglobulin A antibody directed against serotype-specific epitope of Shigella flexneri lipopolysaccharide protects against murine experimental shigellosis. J Exp Medicine 182, 769–778 (1995).

27. S. Boullier, et al., Secretory IgA-mediated neutralization of Shigella flexneri prevents intestinal tissue destruction by down-regulating inflammatory circuits. J Immunol Baltim Md 1950 183, 5879–85 (2009).

28. A. Zychlinsky, M. C. Prevost, P. J. Sansonetti, Shigella flexneri induces apoptosis in infected macrophages. Nature 358, 167–169 (1992).

29. K. Nothelfer, et al., B lymphocytes undergo TLR2-dependent apoptosis upon Shigella infection. J Exp Medicine 211, 1215–29 (2014).

30. W. Salgado-Pabón, et al., Shigella impairs T lymphocyte dynamics in vivo. P Natl Acad Sci Usa 110, 4458–63 (2013).

31. C. Konradt, et al., The Shigella flexneri type three secretion system effector IpgD inhibits T cell migration by manipulating host phosphoinositide metabolism. Cell Host Microbe 9, 263–72 (2011).

32. M. C. Anderson, P. Vonaesch, A. Saffarian, B. S. Marteyn, P. J. Sansonetti, Shigella sonnei Encodes a Functional T6SS Used for Interbacterial Competition and Niche Occupancy. Cell Host Microbe 21, 769–776.e3 (2017).

33. V. Monceaux, et al., Anoxia and glucose supplementation preserve neutrophil viability and function. Blood 128, 993–1002 (2016).

34. P. S. Mitchell, et al., NAIP-NLRC4-deficient mice are susceptible to shigellosis. Biorxiv, 2020.05.16.099929 (2020).

35. B. Marteyn, et al., Modulation of Shigella virulence in response to available oxygen in vivo. Nature 465, 355–358 (2010).

