## Supplementary informations for "The ascorbate-deficient guinea pig model of shigellosis allows the study of the entire *Shigella* life cycle"

**This PDF file includes:**

Figures S1 to S6
SI References

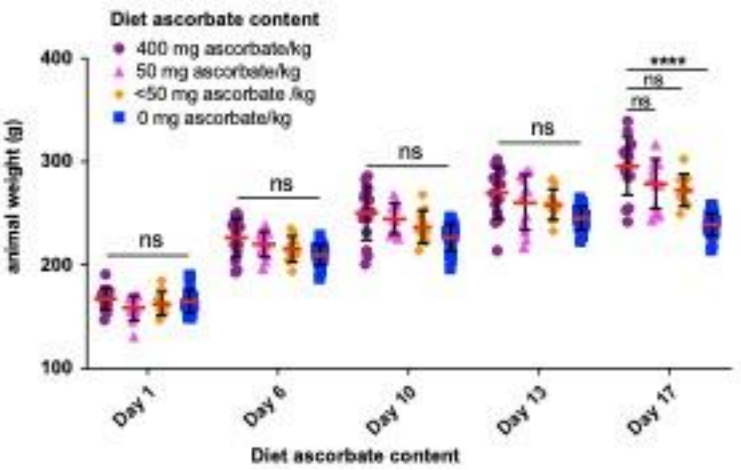


**Figure S1. Impact of diet ascorbate content on guinea pig weight.** Young guinea pigs (3-week) were fed for two weeks with diets containing various ascorbate concentration to induce moderate ascorbate-deficiency without inducing scurvy, caused by severe ascorbate deficiency. Commercial diet contains 400 mg ascorbate/kg (Safediet); suboptimal ascorbate supply was obtained with diets containing 50 mg, <50 mg and 0 mg ascorbate/kg. Animal weight was not significantly different over the 2-week period of time with low-ascorbate containing diets (50 or <50 mg ascorbate/kg diet), it is with a diet devoid of ascorbate (0 mg ascorbate/kg). Results are expressed as Mean ± S.D., **** indicates T-test $p<0.0001$ ($n>10$ animals per group).

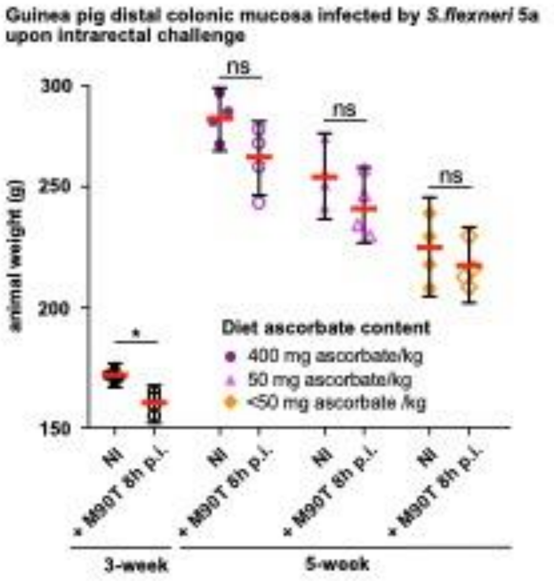


**Figure S2. Impact of 8h Shigella infection on guinea pig weight upon intrarectal challenge.** Young (3-week) and older guinea pigs fed with 400, 50 and <50 mg ascorbate/kg diets (5-week) were challenged intrarectally with $10^{10}$ c.f.u. *S. flexneri* 5a (M90T). 8h p.i. or in the absence of infection (NI, non-infected) animals were weighted. Results are expressed as Mean ± S.D., ‘ns’ indicates T-test *p*>0.05, * indicates *p*<0.05 (4 animals per group).

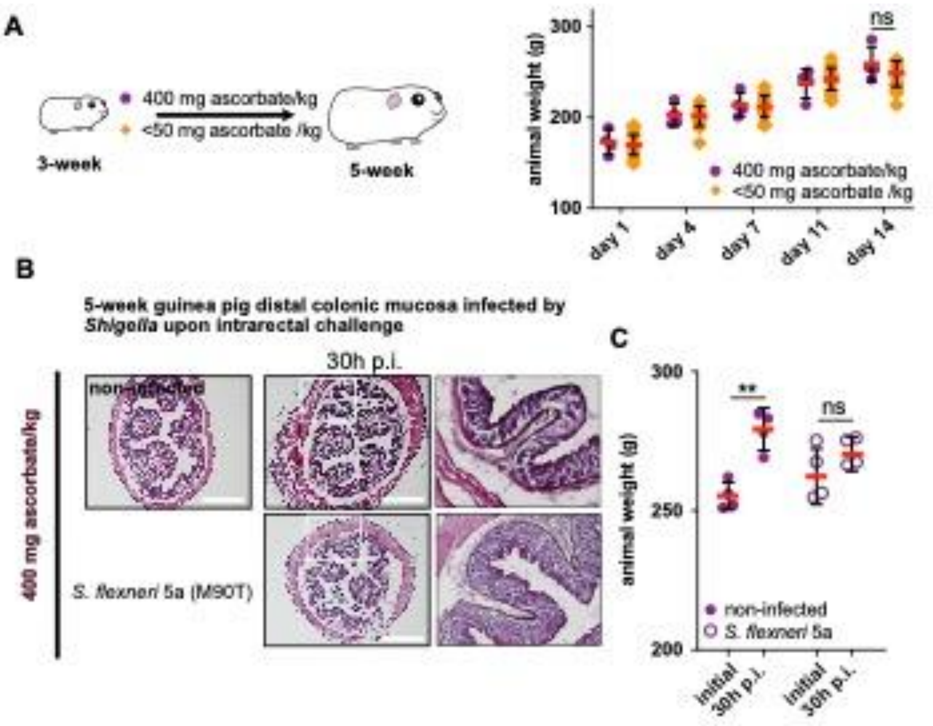


**Figure S3. *Shigella* spp. do not colonize the colonic mucosa of control animals fed with a high-ascorbate containing diet. (A)** Schematic representation of the 2-week moderate ascorbate deficiency induction with a low-ascorbate containing diet (<50 mg ascorbate/kg). Animal weight was recorded twice a week during the ascorbate-deficiency induction. Results are expressed as Mean ± S.D., 'ns' indicates T-test $p$>0.05, (>10 animals per group). (**B**) Ascorbate-deficient guinea pigs (<50 mg ascorbate/kg diet) were infected intrarectally with $10_{10}$ c.f.u. *S. flexneri* 5a (wt). 30h p.i. animals were sacrificed, and distal colonic samples were collected, stained with haematoxylin eosin and compared to non-infected tissues. Scale bars are 30 mm. On the right panel, x3 magnification of white squares (left panel) are shown. (**C**) Animals were weighted before infection and 30h p.i.. Results are expressed as Mean ± S.D., 'ns' indicates T-test $p$>0.05, ** indicates $p$<0.01 (4 animals per group).

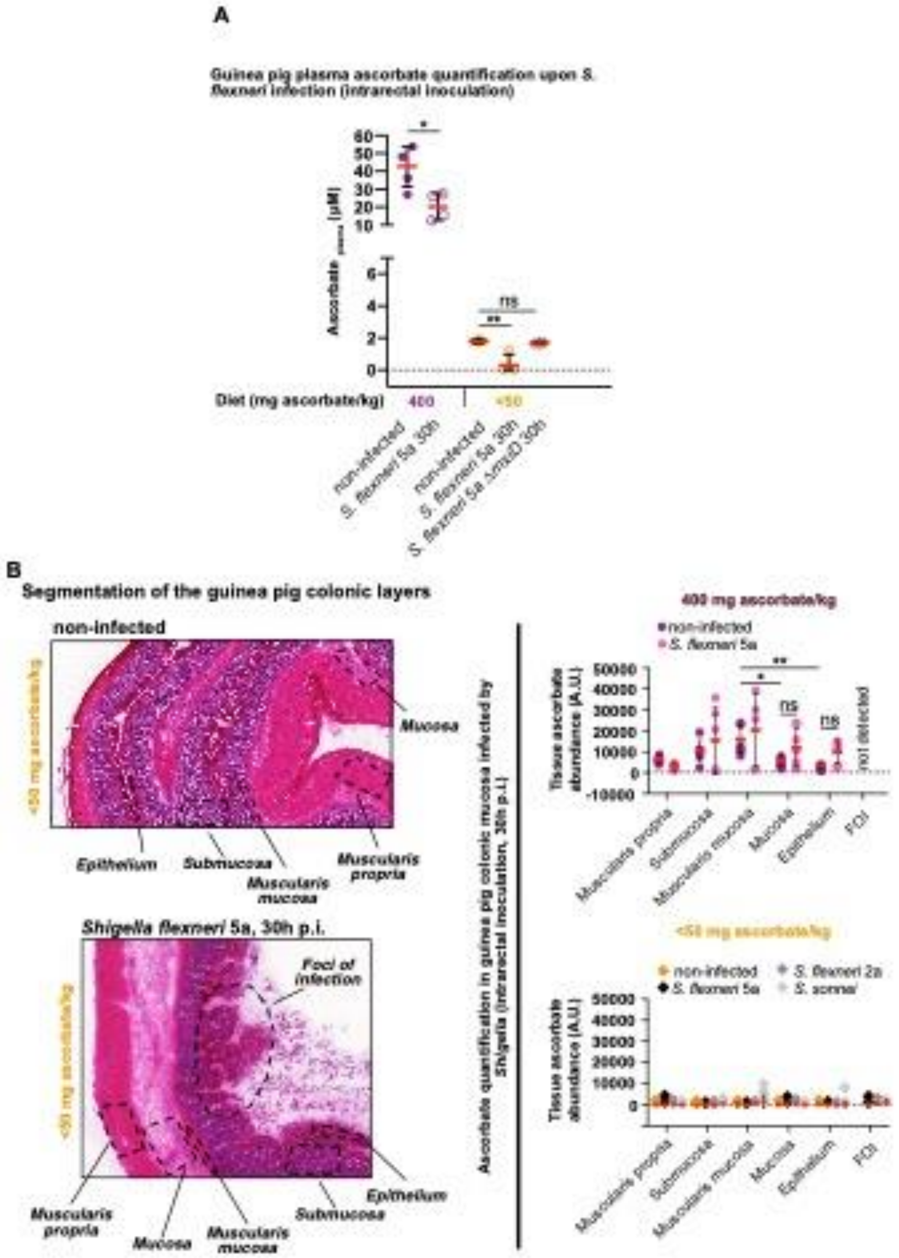


**Figure S4. Plasma and colon ascorbate abundance determination upon *Shigella* infection of ascorbate-deficient and control guinea pigs (A)** Plasma ascorbate concentration (μM) was quantified 30h p.i. in ascorbate-deficient guinea pigs (fed with <50 mg ascorbate/kg diet) and control animals (fed with 400 mg ascorbate/kg diet) upon intrarectal challenge with *S. flexneri* 5a (wt and Δ*mxiD* mutant strains) (**B**) On right panel, the colonic layer segmentation for ascorbate abundance determination is illustrated with two independent samples. On the right panel, ascorbate abundance quantification (imaging mass spectrometry, Arbitrary Units (A.U.)) in the different colon layers from ascorbate-deficient guinea pigs and control animals (400 mg, infected or not by indicated *Shigella* spp. strains (muscularis propria, submucosa, muscularis mucosa, mucosa, epithelium and foci of infection (FOI) when they are detected. Corresponding imaged tissues are shown in Fig. 3C. Results are expressed as Mean ± S.D., 'ns' indicates T-test $p>0.05$, * indicates $p<0.05$, ** indicates $p<0.01$ (n=4 independent areas per group).

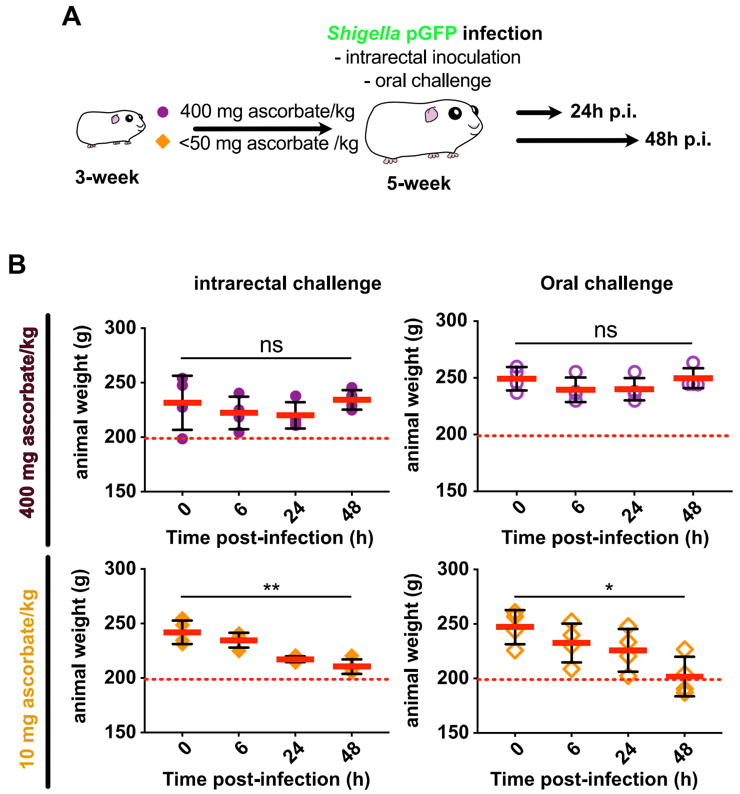


**Figure S5. Plasma and colon ascorbate abundance determination upon *Shigella* infection of ascorbate-deficient and control guinea pigs (A)** Schematic representation of the 2-week moderate ascorbate deficiency induction with a low-ascorbate containing diet (<50 mg ascorbate/kg); followed by an intrarectal or an oral challenge with *S. flexneri* 5a ($10_{10}$ c.f.u.). Animals were sacrificed 24h or 48h p.i. (long-term infections). (**B**) Animal weight was recorded 6h, 24h and 48h p.i.. Experiment was stopped when a 20% reduction weight was reached (dashed red lines). Results are expressed as Mean ± S.D., 'ns' indicates T-test $p$>0.05, * indicates $p$<0.05, ** indicates $p$<0.01 (4 animals per group).

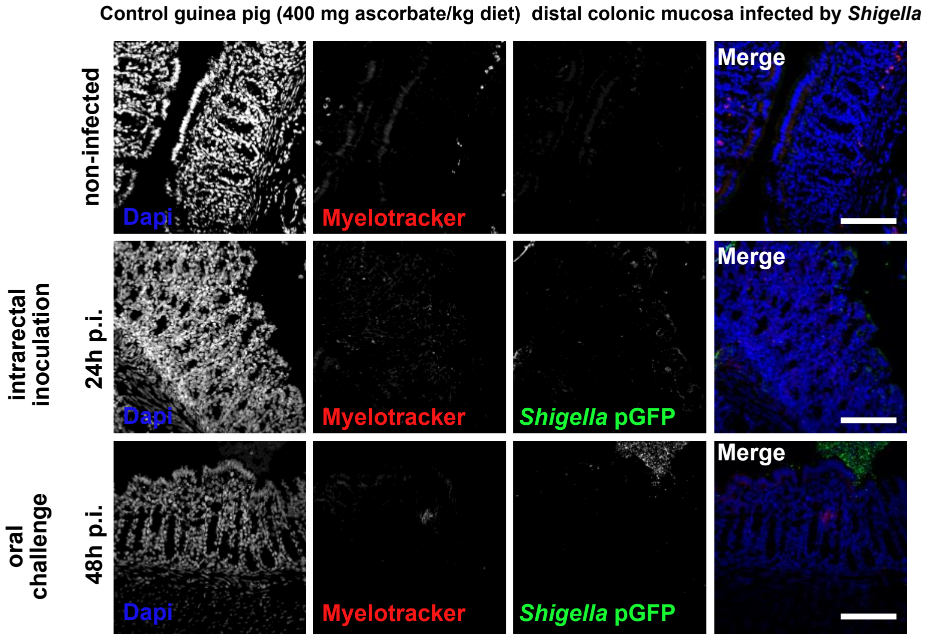


**Figure S6. Shigella do not colonize the colonic mucosa of control guinea pigs (fed with a high-ascorbate containing diet) over extended periods of time.** *S. flexneri* 5a pGFP (green) and neutrophils were detected by immunofluorescence within the distal colonic mucosa of control guinea pigs (400 mg ascorbate/kg diet) challenged orally or intrarectally for 24 and 48h with $10_{10}$ c.f.u.. DNA was stained with Dapi (blue), neutrophils were stained with Myelotracker-Cy3 (18) (red). Scale bars are 150 μm.

**SI References**


1. J.-M. Neuhaus, L. Sticher, F. Meins, Jr., T. Boller, A short C-terminal sequence is necessary and sufficient for the targeting of chitinases to the plant vacuole. *Proc. Natl. Acad. Sci. U.S.A.* 88, 10362–10366 (1991).
2. E. van Sebille, M. Doblin, Data from "Drift in ocean currents impacts intergenerational microbial exposure to temperature." Figshare. Available at https://dx.doi.org/10.6084/m9.figshare.3178534.v2. Deposited 15 April 2016.
3. A. V. S. Hill, "HLA associations with malaria in Africa: Some implications for MHC evolution" in Molecular Evolution of the Major Histocompatibility Complex, J. Klein, D. Klein, Eds. (Springer, 1991), pp. 403–420.
